# DLPFC controls the rapid neural response to visual threat: An ERP and rTMS study

**DOI:** 10.1101/2021.10.04.463069

**Authors:** Justine Cinq-Mars, Anna Blumenthal, Alessa Grund, Sébastien Hétu, Isabelle Blanchette

**Affiliations:** CogNAC research group (Cognition, neuroscience, affect et comportement); Université du Québec à Trois-Rivières, Trois-Rivières, Québec, Canada; Université de Montréal, Montréal, Québec, Canada; Université Laval, Québec, Québec, Canada; Technical University of Munich, Munich, Germany; CERVO Brain Research Center

**Author notes:** Correspondance should be addressed to Isabelle Blanchette, Université Laval, Québec, Canada, Pavillon Félix-Antoine-Savard Local 1144.

**Keywords:** threat superiority effect, visual search task, right DLPFC, rTMS/TMS, EEG/ERP

## Abstract

Individuals are faster at detecting threatening stimuli than neutral stimuli. While generally considered a rapid bottom-up response, this threat superiority effect is also modulated by top-down mechanisms known to rely on the dorsolateral prefrontal cortex (DLPFC). What remains unclear is whether the response is modulated only at later stages of processing, or whether rapid attention to threat *itself* is controlled in a top-down manner. To test this, we used repetitive transcranial magnetic stimulation (rTMS) to inhibit activity in the DLPFC, and measured EEG to index the immediate neural response to threat. Participants attended two sessions where they performed a visual search task with threatening or neutral targets. Prior to this, they received 15 minutes of 1 Hz inhibitory or sham rTMS targeting the right DLPFC. We measured the impact of rTMS on the P1, a rapid visually-evoked potential that is modulated by attention. We found that threatening targets increased the amplitude of the P1 in the sham condition, but inhibition of the DLPFC abolished this increase. These results suggest that the neural signature of rapid attentional detection of threat, even at its earliest stage, is influenced in a top-down fashion by the right DLPFC.

## Introduction

A large body of research has shown that humans orient their attention more rapidly to threatening stimuli than non-threatening stimuli, a phenomenon often called the threat superiority effect (Blanchette, 2006; Brown, El-Deredy, & Blanchette, 2010; Fox, 1996; Fox et al., 2000; Fox, Griggs, & Mouchlianitis, 2007; LoBue, 2014; Mogg & Bradley, 1999; Öhman & Mineka, 2003). For instance, in visual search tasks, snakes and spiders are detected more quickly than benign stimuli (Öhman, Flykt, & Esteves, 2001; Öhman, Lundqvist, & Esteves, 2001). The dominant explanation for this finding is that humans evolved a “fear module’’ for threats common at the time the mammalian brain evolved (Öhman et al., 2001). Specifically, this module has been proposed as a bottom-up circuit, funneling visual information directly to the amygdala. According to this view, amygdala activity increases in response to threatening stimuli, which modulates processing in early visual cortex, leading to rapid attentional-orienting to the threat (Shackman & Fox, 2016).

However, a number of lines of converging evidence complicate this view. First, individuals respond just as rapidly or faster to modern threats, such as syringes and knives, for which learning or experience is necessary, as they do to evolutionarily-relevant threats, such as snakes and spiders, which should evoke the fear module (Blanchette, 2006; Brosch & Sharma, 2005; Fox, Griggs & Mouchlianitis, 2007; Brown et al., 2010; LoBue, 2010; LoBue & DeLoache, 2008). Further, children show this effect specifically for modern threats they have had experience with, such as syringes (LoBue, 2010). Importantly, this enhanced attentional response to modern threats occurs just as rapidly at the neural level as the response to evolutionarily-relevant threats, suggesting a shared neural mechanism (Brown et al., 2010). This can be examined via the P1, a rapid (80-110 ms) parieto-occipital visual evoked potential, thought to be one of the earliest signatures of visual processing modulated by attention (Clark & Hillyard, 1996; Hillyard & Anllo-Vento, 1998; Mangun, 1995; Mangun & Buck, 1998; Luck et al., 2000). Specifically, the amplitude of the P1 is enhanced for both modern and evolutionarily-relevant threatening stimuli compared to neutral stimuli (Brown, El-Deredy, & Blanchette, 2010). In fact, Brown and colleagues (2010) showed that modern threats elicited greater P1 amplitude than evolutionary-relevant ones, corresponding to the finding that individuals were also faster at detecting modern threats.

In addition to the evidence that experience shapes threat detection, there are a growing number of documented top-down influences on the threat superiority effect. For example, strategic processing, based for instance on knowledge about the upcoming stimulus (e.g., you are about to see a snake), facilitates threat detection (LoBue, 2014). Detection is also facilitated when individuals are instructed to use an emotional processing strategy, as opposed to a semantic one (being asked if the stimulus is dangerous vs. being asked if it is an object; Damjanovic, Williot & Blanchette, 2020). The dorsolateral prefrontal cortex (DLPFC) likely underlies this response modulation (Ochsner & Gross, 2005). Indeed, DLPFC is involved in categorization and evaluation of emotional stimuli (Zwanzger et al., 2014), and plays a causal role in top-down modulation of visual attention more generally (Noudoost, Chang, Steinmetz, & Moore, 2010; Sagliano et al., 2016; Zanto et al., 2011). Further, DLPFC has direct projections to posterior visual areas, making direct rapid modulation of early visual attention possible (Pierrot□Deseilligny, Müri, Nyffeler, & Milea, 2005). This raises the interesting possibility that the DLPFC may modulate the initial rapid attentional orientation to threat. If the DLPFC directly controls rapid attention, an increase in DLPFC activity may be required to increase the attentional processing of threatening stimuli, as reflected in the increase in amplitude in the P1 to threat. If this is the case, inhibiting the DLPFC should reduce the normal increase in P1 amplitude to threatening stimuli and corresponding reaction times should be slower. This prediction is consistent with the DLPFC connectivity with visual cortex.

This prediction is in stark contrast to the hypothesized role of the DLPFC according to the *emotion-regulation theory*. According to this theory, the role of the DLPFC is essentially one of downregulation. Rapid attention to threat occurs through amygdala activation which directly modulates visual processing. This is supported by the finding that patients with amygdala lesions do not show increased BOLD activity in the visual cortex in response to threat (Vuilleumier et al., 2004). The DLPFC is thought to inhibit amygdala activity when necessary (De Raedt et al., 2010). Hence, decreased DLPFC activity should correspond to increased amygdala activity, with greater visual evoked responses and faster reaction times toward threatening stimuli.

To evaluate whether the DLPFC plays a causal role in rapid attentional orienting and delineate between these theories, we used repetitive transcranial magnetic stimulation (rTMS) combined with EEG.

rTMS can be used to selectively excite or inhibit the DLPFC through high or low-frequency stimulation (Sliwinska, Vitello, & Devlin, 2014), and can modulate behavioral responses to threat (De Raedt et al., 2010; Sagliano et al., 2016; Zwanzger et al., 2014). One previous study found that inhibition of the right DLPFC led to slower reaction times categorizing fearful faces (Zwanzger et al., 2014), while another found that excitation of the right DLPFC led to slower attentional disengagement from angry faces (De Raedt et al., 2010). Both of these studies show the possible involvement of the DLPFC in processing negative, potentially threatening stimuli, but they both use complex tasks that involve strategic processes in addition to initial detection. This leaves open the question of the potential involvement of the DLPFC in modulating immediate responses to threat.

Thus, to our knowledge, there are no studies using a visual search threat detection task to investigate the role of the DLPFC in modulating the initial threat detection response. This is an important step forward, as the visual search task is the main paradigm used to study the rapidity, or efficiency, with which people detect threatening stimuli (Blanchette, 2006; LoBue, 2014; LoBue & DeLoache, 2010; Öhman, Flykt, & Esteves, 2001a; Öhman, Lundqvist, & Esteves, 2001b; Treisman & Gelade, 1980; Treisman & Souther, 1985; Turatto & Galfano, 2004; Wolfe, 1998), it also has ecological validity — it has been linked to clinical issues such as phobia and anxiety (Ohman et al 2001a; LoBue, 2014). Here, we combine inhibitory rTMS with EEG to see if right DLPFC modulates one of the most rapid neural markers of visual attention, the P1.

### Aim of the study and hypothesis

Our aim was to explore the role of the right DLPFC in early neural signature of attention to threat. To do this, participants received inhibitory or sham-rTMS, and then performed a visual search task in which they had to identify targets (threatening or neutral) among distractors (Blanchette, 2006). We expected to replicate the canonical finding of faster detection of threatening stimuli compared to neutral ones, in the sham-rTMS condition (Blanchette, 2006; Brown et al., 2010; Fox et al., 2007; LoBue, 2014; Öhman & Mineka, 2003). Further, we expected that this threat superiority effect would be greater for modern threats, given prior findings with studies including participants with similar demographics (Brown et al., 2010). We also expected that this sensitivity to threat would be reflected in an elevated P1 amplitude for threatening targets compared to neutral ones (Bar-Haim et al., 2005; Carretié et al., 2001; Huang & Luo, 2006; Zwanzger et al., 2014), and that this would be increased for modern threats (Brown et al., 2010).

The novel hypothesis we explore is that inhibiting right DLPFC activity would reduce the threat superiority effect, reflected in slower reaction times to threatening targets in the inhibition rTMS condition. Crucially, we anticipated that this reduction in attention to threat would occur at the earliest stages of neural processing, as evidenced by a reduction in the amplitude of the P1 for threatening targets compared to neutral targets. We expected similar rTMS effects for both evolutionary-relevant and modern threats.

## Materials and methods

### Participants

Twenty individuals (13 females, mean age = 21.8, *SD* = 2.0) participated in this study. They gave written informed consent and received fifty dollars for their participation. All participants had normal or corrected-to-normal vision, were right-handed, had no reported history of psychiatric or neurological disorders, and no contraindications for rTMS stimulation (Rossini et al., 2015). One female participant interrupted her participation in the experiment because of a headache, leaving 19 participants in the final analyses. The local ethics committee of the Université du Québec à Trois-Rivières approved the study, and the rTMS protocol was in agreement with the International Federation of Clinical Neurophysiology safety guidelines (Rossini et al., 2015).

### Apparatus, stimuli, and design

Participants were tested in an isolated 10×10 feet Faraday cage. The experiment was programmed and executed using E-Prime 3.0 software, and displayed via an Intel Core i7 computer, using a 23-inches monitor.

#### TMS

We delivered TMS using a double 70 mm air-cooled figure-eight coil connected to a Magstim Rapid 2 Plus 1 stimulator. We estimated the site of stimulation with the F4 electrode in the 10-20 EEG coordinate system. This method has been used before in rTMS research to inhibit the right DLPFC (Schutter, van Honk, d’Alfonso, Postma, & de Haan, 2001; Rusjan et al., 2010; Zwanzger et al., 2014). Inhibition intensity was set at 120% resting motor threshold (MT) but was reduced to 100% for nine participants who reported headaches at the beginning of the train of pulses. Reduction of intensity is a common practice in rTMS studies (Maizey et al., 2013). There was no link between the stimulation’s intensity and the main effects of interest (relative P1 amplitude to threat and neutral stimuli) [*r* = -0.04, *p* = 0.86].

#### EEG recording

We recorded the continuous EEG signal using a TrueScan 7.5 recording system and tin electrodes compatible with TMS (Deymed Diagnostic, Czech Republic). Sixty-four channels were recorded using the International 10/20 System sites. A reference montage was chosen with CZ as a reference and the right mastoid as the ground. The impedance was kept under 20 ohms for all electrodes. The signal was digitalized at 3000 Hz.

### Visual Search Task

The visual search task was based on a previous study (Blanchette, 2006). The display consisted of a matrix of nine greyscales photos equally spaced in a rectangle shape on a white background. There were two types of matrices. The first type consisted of eight images from the same category—the distractors—and one target image from a different one (e.g., a spider with palm trees). The second type of matrix presented only images of the same category (e.g., only spiders or only palm trees). Examples are presented in Figure 1. The first type was presented for two-thirds of trials (200 times) and the second type for one third (100 times).

**Figure 1.**
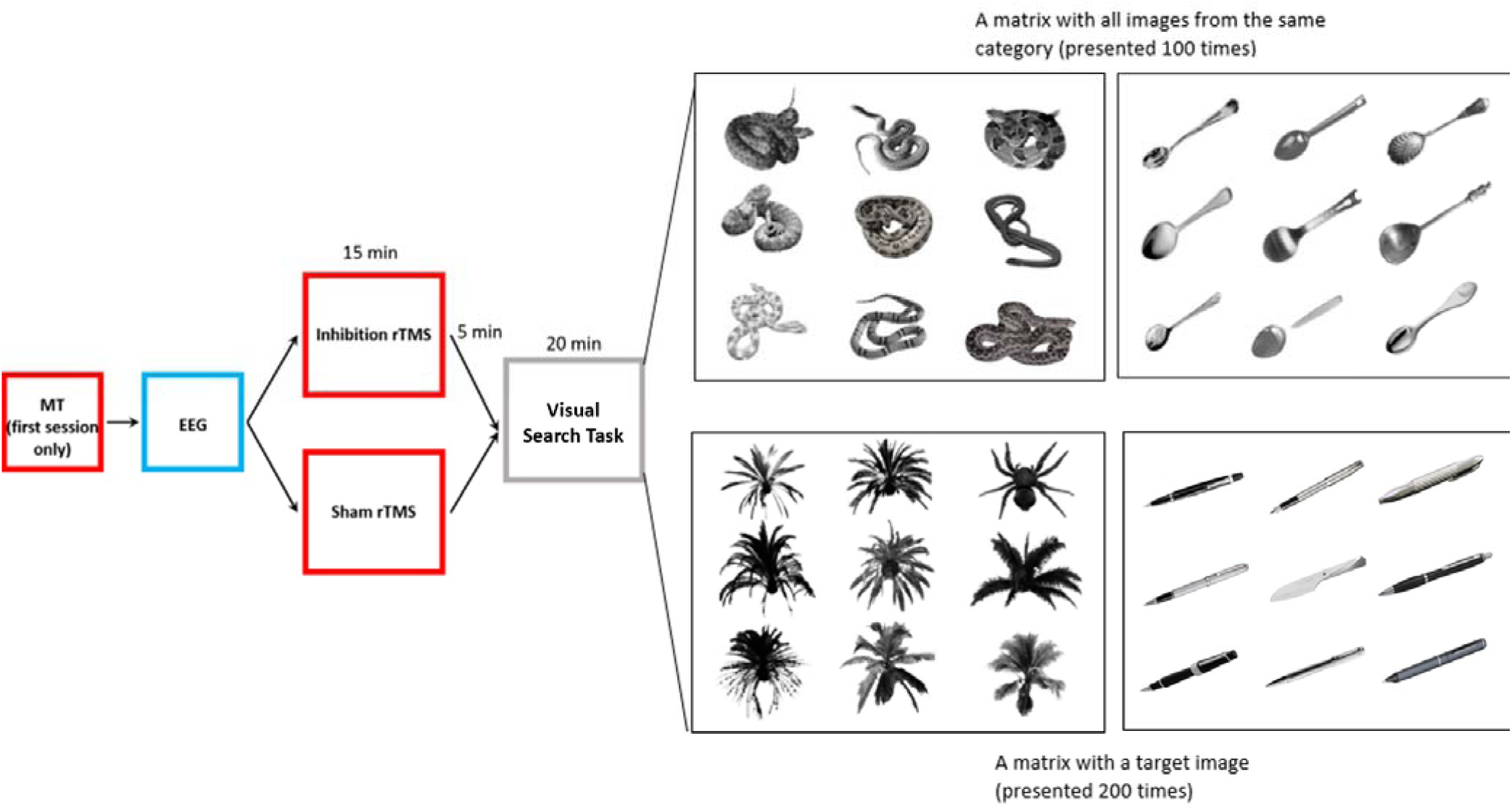
Experiment Procedure

Targets varied along two dimensions: threat-relevance and evolutionary relevance. Manipulation of threat-relevance and evolutionary relevance created four types of matrices (see Table 1).

**Table 1.** All types of targets presented, by threat and evolutionary relevance (with their distractors)

|  | Evolutionary-relevant | Evolutionary-irrelevant |
| --- | --- | --- |
| **Threat** | Snake (rope) | Gun (spoon) |
|  | Spider (palm tree) | Knife (pen) |
| **Neutral** | Rope (snake) | Spoon (gun) |
|  | Palm tree (spider) | Pen (knife) |

Participants were instructed to determine whether all objects were from the same category or whether there was one object from a different category. They were asked to respond as quickly and as accurately as possible by pressing response keys (‘A’ or ‘L’) with their index fingers. Response key correspondence was counterbalanced across participants. They were also instructed to keep their eyes on the fixation cross and avoid eye movements and blinks during matrix presentations. The task began with eleven practice trials, including feedback on accuracy. Altogether, 300 trials were presented in random order without feedback. The position of the target varied randomly across matrices. Each trial started with a black fixation cross in the center of the screen, presented between 600 to 1100 ms, randomly. Three one-minute breaks were offered during the task every 100 trials.

### Experimental procedure

Participants took part in two sessions, conducted on different days (at least one week apart): an inhibition rTMS condition and a sham rTMS condition. Condition order varied randomly between participants. During the first session, the resting motor threshold (MT) was determined by the thumb movement visualization method with a single pulse TMS (Rossini et al., 2015). This represents the minimum pulse intensity required to produce five muscle contractions (>0.5 mV) of the abductor brevis muscle of the thumb during ten consecutive stimulations. The NDI Polaris Vicra Camera neuronavigation system was used to position the coil, and muscle contractions were measured using a CED (Cambridge Electronic Design, UK) 1902 isolated dual channel preamplifier with EMG adapter box, digitized by a CED Micro1401-3 data acquisition unit and Spike 2 software. After that, both sessions followed the same procedure (see Figure 1). Participants were seated 70 cm away from the screen where they would perform the visual search task. The EEG cap was first installed. Participants then received 1-Hz train of pulses with the rTMS (inhibition or sham) for 15 minutes (900 pulses) on their right DLPFC. The effects of 1-Hz stimulation last approximately 15 to 20 minutes (Fitzgerald & Daskalakis, 2013). In the sham rTMS condition, the coil position was rotated 90 degrees, so only one wing touched the F4 electrode. This type of sham rTMS produces no to minimal stimulation (Loo et al., 2000; Valero-Cabré et al., 2011). Afterward, participants performed the visual search task.

The time between the end of the rTMS stimulation and the beginning of the visual search task was approximately five minutes (impedance of electrodes was verified and adjusted). The task required approximately 20 minutes to complete. Because this sequence exceeded the inhibition time effect of 15-20 min, we conducted analyses to examine if there was evidence of the effect fading with time. These analyses are reported in the results section and confirm a main effect of time, but that did not vary between rTMS conditions.

### Behavioral data analysis

Accuracy was measured to be sure no condition was harder than any other. Accuracy did not significantly differ across rTMS conditions and threat relevance of the target [*F*(1,18) = 0.38, *p* = 0.54, η_p_^2^ = 0.02]: inhibition threat (*M* = 93.3%, *SD* = 1.5); inhibition neutral (*M* = 89.1%, *SD* = 2.1); sham threat (*M* = 94.6%, *SD* = 1.0); sham neutral (*M* = 89.8.3%, *SD* = 2.0). Our behavioral dependent measure was reaction time (RT). For RT analyses, we only included accurate target-present trials. We also excluded trials with RTs <500 ms or >3000 ms, resulting in the exclusion of 2,5% of trials. We conducted one repeated-measures ANOVA on RTs with the following factors: rTMS condition (inhibition or sham), threat relevance of the target (threatening or neutral), and evolutionary relevance of the target (evolutionary-relevant or modern)

### EEG data analysis

We performed EEG signal processing and analysis using Brain Vision Analyzer 2.0 software (Brain Products, Gilching, Germany). The EEG data were high-pass filtered with a cutoff of 0.01 Hz and low-pass filtered with a cutoff of 80 Hz (IIR filter). Filtered data was segmented for each 200 trials with a target present, beginning 200 ms before matrix presentation until 600 ms after stimulus onset (for a total of 800 ms). The 100 target-absent trials (all the same category stimuli) were not use for analysis, as they were only created to make the task possible.

Baseline correction was performed using the period from 200 ms to 25 ms prior stimulus onset. This was used to avoid an electric artifact appearing between 10 ms before stimulus onset and offsetting 10 ms after stimulus presentation. This artifact was due to a broken cable, it stopped when the cable was changed after the experiment, and did not impact the component of interest (it was concise and was outside the P1 time window). We rejected segments containing artifacts detected by a voltage difference higher than 80 μV within 100 ms intervals on channels of interest (PO7 and PO8, see below). This criterion led to a rejection of 4,9% trials among all participants (varying from: 1,3% of rejected trials to 8,9% between participants)

Time window of P1 was defined as the mean amplitude from 75 to 125 ms after target onset. This time window was chosen from inspecting the grand average waveform (averaged over all participants and conditions), showing a clear P1 at 100 ms post-stimulus (see Figure 3). We localized P1 at parieto-occipital sites, where the P1 is typically maximal (Luck et al., 2000). Specifically, electrodes PO7 and PO8 were chosen as the electrodes of interest based on data inspection. Also, PO7 and PO8 were collapsed to facilitate the analysis, as P1 is maximal on the hemisphere contralateral to stimulus presentation and the position of the target varied randomly from left visual field to right visual field during the task.

Repeated measures ANOVAs were conducted to investigate the main effect of rTMS condition and target type on the amplitude of the P1. To follow up on significant interactions, we conducted paired-sample t-tests with Bonferroni correction. We estimated partial eta-squared (η_p_^2^) and Cohen’s *d* to describe effect sizes. Based on Cohen’s guidelines, η_p_^2^ values from 0.01 to 0.05 reflect a small effect, values from 0.06 to 0.13 indicate a medium effect, and values ≥ 0.14 indicate a large effect; and *d* values from 0.2 to 0.4 indicate a small effect, value from 0.5 to 0.7 a medium effect and values ≥ 0.8 indicate a large effect (Cohen, 1988).

## Results

### Reaction times

Mean RTs to detect targets are shown in Figure 2. Overall, participants detected threatening targets faster than neutral targets, [*F*(1,18) = 25.78, *p* □ 0.001, η_p_^2^ = 0.59]. They also detected modern targets faster than evolutionary-relevant ones [*F*(1,18) = 68.42, *p* □ 0.001, η_p_^2^ = 0.79]. No main effect of rTMS condition was found [*F*(1,18) = 0.17, *p* = 0.69, η_p_^2^ = 0.009], and no interaction of rTMS condition with threat relevance of the target [*F*(1,18) = 0.45, *p* = 0.53, η_p_^2^ = 0.03] or with evolutionary-relevance were found [*F*(1,18) = 0.22, *p* = 0.65, η_p_^2^ = 0.01], nor interaction of threat relevance of the target with evolutionary-relevance [*F*(1,18) = 3.35, *p* = 0.08, η_p_^2^ = 0.16].

**Figure 2.**
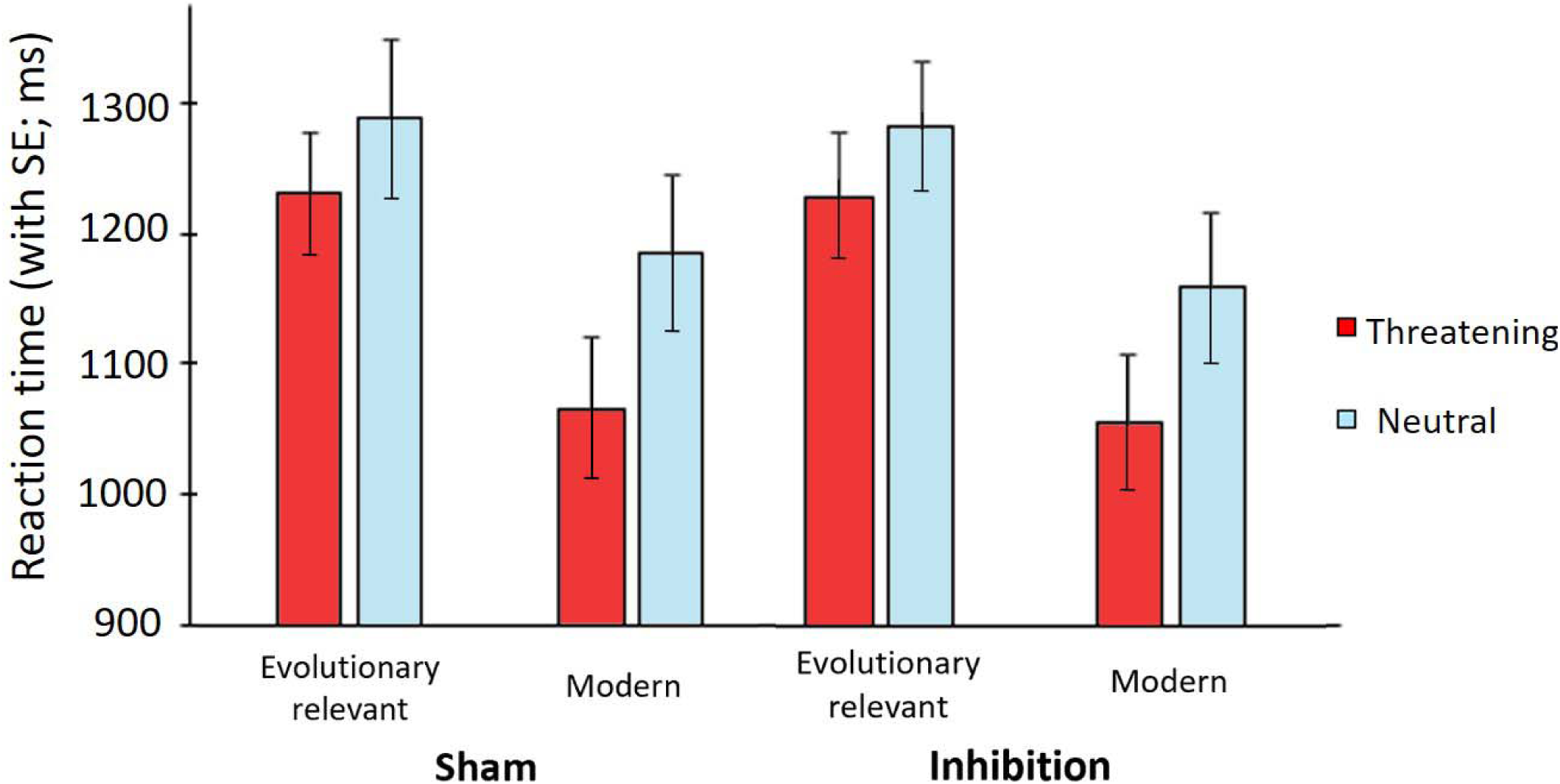
Reaction Time

### Effect of time

As mentioned before, the preparation for the task and the task itself exceeded the expected length of the inhibition, typically estimated at 15-20 min. Hence, we conducted analyses to determine whether the effect of rTMS stimulation diminished with time. We conducted a one-way ANOVA with the following factors: time (first 100 target-present trials vs. last 100), threat relevance of the target (threatening vs. neutral), and rTMS condition. There was no time x rTMS condition interaction on reaction times [*F*(1,15) = 0.12, *p* = 0.74, η ^2^ = 0.008]. However, a main effect of time was observed, with longer reaction time for the last trials, (a commonly seen effect related to fatigue).

### ERP waveforms

Grand-average waveforms time-locked to stimulus onset, collapsed across evolutionary relevance and across PO7 and PO8 electrodes sites are presented in Figure 3.

**Figure 3a.**
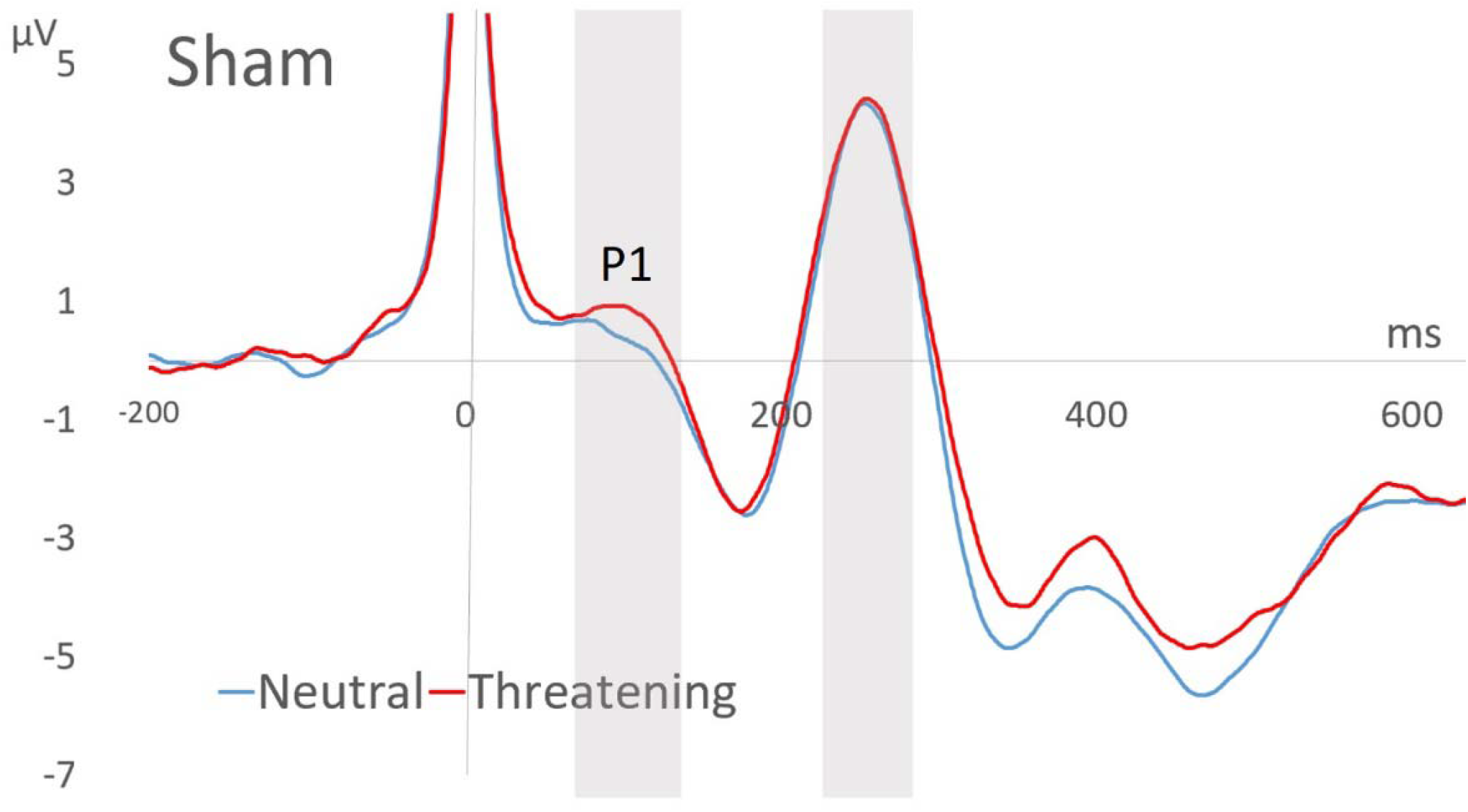
ERP Waveform (Sham)

**Figure 3b.**
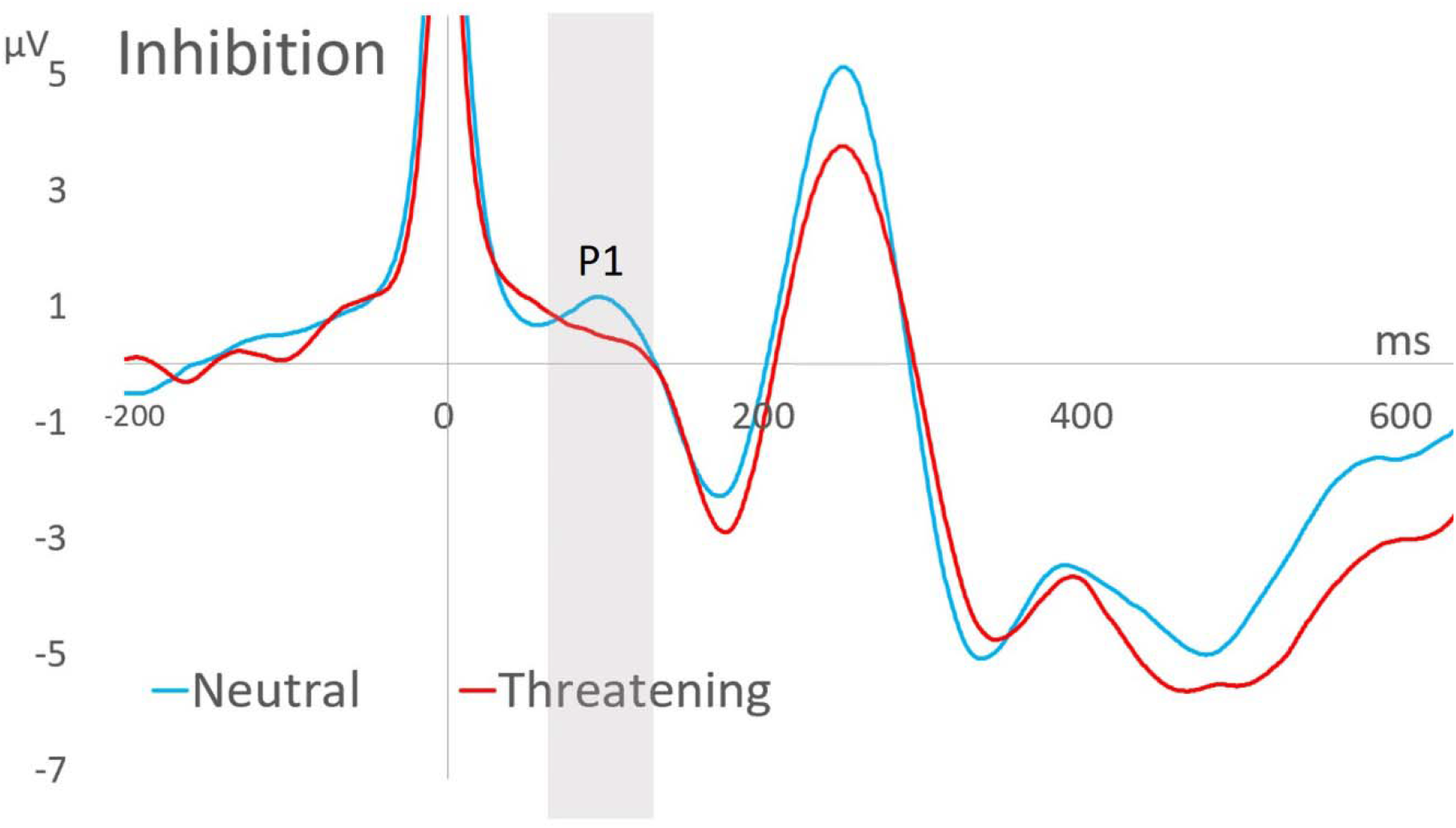
ERP Waveform (DLPFC Inhibition)

**Figure 3.** Grand average of P1 by rTMS condition (a. sham, b. inhibition) and threat relevance of the target (red = threatening; blue= neutral). Zero ms corresponds to stimulus presentation. The cable artifact can be seen around 0 ms but did not impact analysis, as mentioned in the Materials and methods section.

#### P1 component

The ANOVA revealed a significant interaction effect between rTMS condition and threat relevance of the target on P1 amplitude [*F*(1,18) = 5.73, *p* = 0.03, η_p_^2^ = 0.24; see Figure 4]. In the sham-rTMS condition, the amplitude of the P1 tended to be larger for threatening targets (*M* = 2.63, *SE* = 0.51) compared to neutral ones (*M* = 1.58, *SE* = 0.40), with a strong effect size [*t*(18) = 2.0, *p* = 0.12, *d* = 2.3, two-tailed]. However, in the inhibition rTMS condition, P1 amplitude was equivalent for threatening and neutral targets [*t*(18) = 0.07, *p* = 0.95, two-tailed]. Other interactions and main effects were not significant.

**Figure 4.**
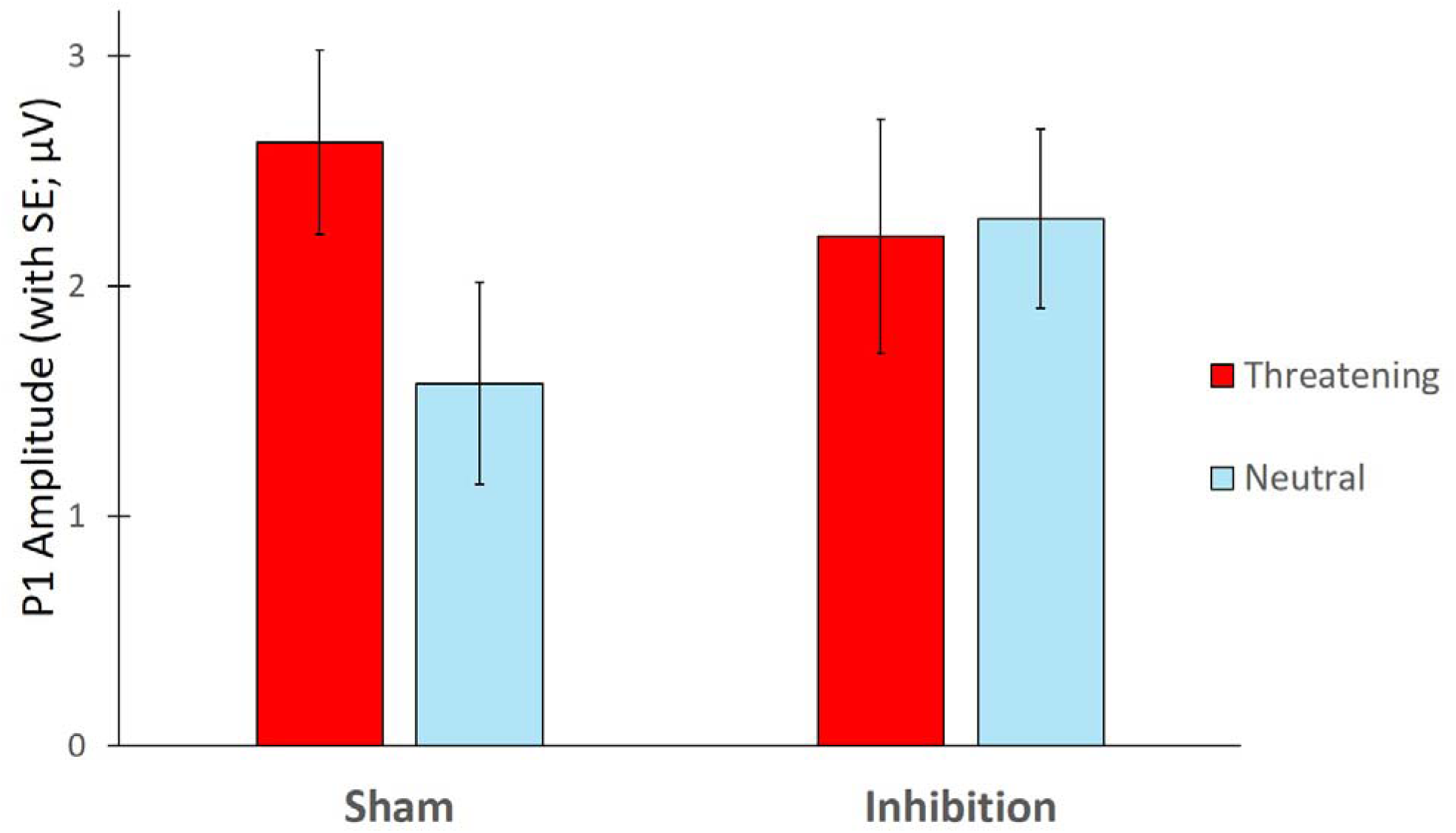
Amplitude P1

An interaction was found between threat relevance and evolutionary-relevance of the target [*F*(1,18) = 5.37, *p* = 0.03], indicating that among modern stimuli, threatening targets elicited larger P1 amplitude (*M* = 2.90, *SE* = 0.47) compared to modern neutral targets (*M* = 1.78, *SE* = 0.42; *t* (18) = -2,42, *p* = 0.52), which was not seen among evolutionary-relevant ones (threatening: *M* = 1.94, *SE* = 0.45; neutral: *M* = 2.09, *SE* = 0.38; *t* (18) = -0.76, *p* = 0.46). This replicates results found in a previous study (Brown et al., 2010). Evolutionary relevance did not, however, interact with rTMS condition [*F*(1,18) = 0.22, *p* = 0.65]. Therefore, the results above are reported collapsed across evolutionary-relevant and modern threats.

## Discussion

We used neuromodulation in conjunction with EEG to test the hypothesis that rapid neural signature of attention to threat is orchestrated top-down by the DLPFC. Crucially, in support of our hypothesis, we found that inhibition of the right DLPFC abolishes the increase in one of the most rapid neural signals of attention to threat – the P1. Specifically, our main findings can be summarized as follows, 1) threatening stimuli were detected more quickly than neutral stimuli, replicating the canonical threat superiority effect, 2) threatening targets enhanced the amplitude of the P1 visually-evoked potential relative to neutral targets, confirming prior findings on the neural markers of attention to threat, and, crucially, 3) inhibition of the right DLPFC modulated this neural marker of the threat superiority effect, suggesting an important role for this structure in early attention towards threatening stimuli.

Our key finding, in support of our hypothesis about the role of the DLPFC in early attentional modulation, is that rTMS stimulation of the right DLPFC modulated the variation in P1 amplitude as a function of threat. Specifically, in our sham condition, we found evidence of rapid neural modulation by the presence of threat - the P1 amplitude was larger for threatening targets, in line with existing literature (e.g., Blanchette, 2006; Brown et al., 2010; Fox et al., 2007; LoBue, 2014; Öhman & Mineka, 2003). Interestingly, participants were faster at detecting modern threats, and there was also an increase in P1 amplitude for modern threats, replicating the findings in Brown et al. (2010). This increase in P1 amplitude to threatening stimuli across all threat types was abolished by rTMS inhibition to the right DLPFC.

These results indicate that DLPFC controls the neural marker of attentional selection to threat at the most rapid timescale/earliest stages, challenging the idea that the threat superiority effect is only a bottom-up response, and that DLPFC’s role in this effect is as an inhibitor of amygdala activity. Indeed, according to the emotional-regulation model, inhibiting DLPFC should lead to an enhanced amygdala response and a stronger threat superiority effect (Bishop et al., 2004; Fales et al., 2008; Peers et al., 2013.), not the reduced response we see. Our results, in contrast, support the hypothesis that the right DLPFC is involved in a direct and fast attentional selection toward threat (De Raedt et al., 2010). This is in line with previous work that showed inhibition of the DLPFC reduced attention to threat behaviorally (Zwanzger et al., 2014). Our study also provides the first evidence that inhibition of DLPFC impacts the rapid neural signature of visual attention. The P1 is recorded from electrodes in occipito-parietal cortex (Di Russo et al., 2002; Zwanzger et al., 2014), and is thought to reflect processing in extrastriate visual cortex (Di Russo et al., 2002). Thus finding that the P1 is impacted by DLPFC inhibition demonstrates that the DLPFC can exert a rapid influence on posterior and even distant visual brain areas, in line with its connectivity and with previous work showing it plays a causal role more generally in visual attention (Zanto et al., 2011). Whether this effect occurs through direct connections, or through connectivity between the DLPFC, amygdala, and visual cortex, remains to be tested by future fMRI studies. However, this finding clearly cannot be accounted by a basic model of emotional regulation centered around the amygdala, which predicts the opposite pattern of results. If the DLPFC impacts threat detection via inhibition of the amygdala, it is through a more nuanced and rapid interaction than previously theorized.

Somewhat surprisingly, we did not see a behavioral impact of abolishing the P1 sensitivity to threat with DLPFC inhibition. Specifically, we observed the same robust threat superiority effect in the rTMS condition as the sham condition. Importantly, other studies have shown a similar pattern, indicating that it is possible to observe a shift in the electrophysiological mechanisms involved in threat processing in the absence of a change in behavior (Kappenman et al., 2015; Brown et al., 2010). This is also in accordance with the result of Sagliano et al. (2016), who did not find behavioral results after inhibiting the right DLPFC. This could be explained by the precocity (100-200 ms) of the cognitive processes indexed by the P1. It is possible that other processes could have restored the usual attention allocated to threats subsequently. For example, other regions known to be involved in ongoing attention and emotional response regulation, such as the ventro-lateral prefrontal cortex or anterior cingulate cortex (Pessoa & Adolphs, 2010; Scott et al., 2015) could orient attention to threat at a later stage.

To our knowledge, the present study is the first to combine EEG and rTMS to uncover the neural signature of rapid visual attention to threat. Due to the challenging nature of the study, the sample size (n=19) is relatively modest. However, despite this modest sample size we replicate key behavioral and ERP findings (the threat superiority effect, the increase in the effect for modern threats, the P1 elevation for threats, and the increased elevation for modern threats). Furthermore, we see a strong effect (n_p_^2^ = 0,24) of rTMS inhibiting the right DLPFC on P1 amplitude, specifically abolishing the initial P1 elevation to threat, confirming our hypothesis that top-down mechanisms play an important role in determining responses to threat from an early stage.

## Conclusion

To conclude, neuroimaging combined with neurostimulation allowed us to investigate the early stage of attentional selection toward threats. Considering that the source of P1 component is located in the extrastriate occipito-parietal cortex (Di Russo et al., 2002; Zwanzger et al., 2014), these results demonstrate that the right DLPFC can exert influences on posterior and even distant visual brain areas, at the earliest stages of threat perception. Our study provides new insights into one of the neural mechanisms possibly underpinning the threat superiority effect, and that this core aspect of visual perception, even at the most rapid timescale, is controlled top-down.

## Supplementary data Conflict of interest

None declared

## Acknowledgements

This research was funded by an The Natural Sciences and Engineering Research Council of Canada (NSERC) Discovery grant awarded to IB.

## Notes

### Competing Interest Statement

The authors have declared no competing interest.

